# ΔCt-Informed, Calibrated Logistic Regression Accurately Attributes *mecA* in *Staphylococcus aureus*–Positive Wound Specimens

**DOI:** 10.1101/2025.10.03.680191

**Authors:** Mehdi Dehghani, Hans Norouzi, Shabnam Dehghan, Keagan H. Lee, Howard L. Martin

## Abstract

In wound specimens, co-detection of *mecA* and *Staphylococcus aureus* by PCR does not necessarily indicate MRSA because coagulase-negative *staphylococci* (CoNS) frequently harbor *mecA*. We evaluated a ΔCt-informed, biologically gated, calibrated logistic regression to attribute mecA to *S. aureus* versus CoNS. Using paired culture/AST and multiplex real-time PCR Ct values (internal n=93; external n=47), we trained 5-fold cross-validated models in the culture-positive *S. aureus* subset (n=36) and applied an *S. aureus* PCR gate (no attribution when *S. aureus* PCR is negative). The primary model achieved sensitivity 90.9% and specificity 92.0% for MRSA attribution with AUC 0.931 (out-of-fold). Decision curve analysis showed positive net benefit across clinically relevant thresholds; at the prespecified 50% cutoff, the model achieved a net benefit of 0.222 compared with negative benefit for a treat-all strategy. In an external cohort, *S. aureus* detection by PCR versus culture showed 92.3% sensitivity and 97.1% specificity; within *S. aureus* PCR-positives (n=12), MRSA attribution reached 100% sensitivity and 87.5% specificity (accuracy = 91.7%). This framework improves *mecA* interpretability in polymicrobial specimens and can reduce unnecessary MRSA-directed antibiotics.

## Introduction

Wound infections are often polymicrobial, commonly involving pathogenic *Staphylococcus aureus* and commensal coagulase-negative *staphylococci* (CoNS) (1, 2). *S. aureus* is a major cause of invasive infections, whereas CoNS, particularly *Staphylococcus epidermidis*, frequently colonize chronic wounds and indwelling devices. In these contexts, CoNS contribute to biofilm formation and acts as reservoirs of antibiotic resistance determinants, such as *the mecA* gene (3, 4). The presence of the *mecA* resistance gene complicates diagnostic interpretation in such polymicrobial settings. Because *mecA* encodes penicillin-binding protein 2a (PBP2a), conferring methicillin resistance, and its detection in an *S. aureus*–positive specimen may be interpreted as MRSA. However, CoNS commonly harbor *mecA* on *staphylococcal cassette chromosome mec* (*SCCmec)* elements, so assuming that *mecA* must originate from *S. aureus* can be misleading and can prompt unnecessary use of antibiotics such as vancomycin or linezolid [5], instead of β-lactams (e.g., amoxicillin–clavulanate) when the infection is methicillin-susceptible. Such unwarranted treatment escalations increase drug toxicity, cost, and sometimes complicate treatment strategy (1, 3, 5). These insights underscore the need for diagnostic strategies that explicitly account for polymicrobial samples. To address this issue, previous studies used ΔCt (*S. aureus* Ct – *mecA* Ct) thresholds to determine the connection between these two targets. For instance, the MRSA/SA ELITe MGB kit considers a small difference (ΔCt < 2 cycles) between *mecA* and *S. aureus* signals as indicative of MRSA. In contrast, larger differences imply MSSA or mixed populations (6). Another approach combining selective broth enrichment and real-time nuc–*mecA* duplex PCR achieved a sensitivity of 93.5 % and specificity of 88.6 % for MRSA detection (7).

Although these stand-alone MRSA assays exist and have utility in nasal screening or targeted surveillance, they are not optimized for polymicrobial wound specimen testing. In contrast, many CAP/CLIA-certified diagnostic laboratories now use panel-based, probe-based TaqMan assays offered as laboratory-developed tests (LDTs) to provide comprehensive detection of wound pathogens and resistance genes. Our approach was designed to enhance these multiplex workflows by refining the attribution of *mecA* signals within the panel, rather than relying on a separate single-target assay. By embedding ΔCt-derived features and biologic gating into the panel framework, we improve the interpretability of resistance gene results without altering established laboratory workflows. Overall, these studies indicate the potential of quantitative PCR kinetics to improve MRSA diagnosis. we integrated ΔCt information into a calibrated logistic regression model with biologic gating to attribute *mecA* to *S. aureus* versus CoNS in wound infection specimens.

## Methods

### Study Design and Ethics

This diagnostic accuracy study (Supplementary Figure S1) followed the Standards for Reporting of Diagnostic Accuracy Studies (STARD 2015) guidelines (8). We analyzed an anonymized dataset that was previously utilized in the study titled “Comparative Diagnostic Evaluation of Real-Time PCR and Culture for Detecting Pathogens in Podiatric Wound Infections” (9). The study involved paired aerobic culture and antimicrobial susceptibility testing (AST), conducted by a reputable commercial reference laboratory, along with a probe-based Wound PCR panel (BioExcel Diagnostics). Since the dataset was fully de-identified, the analysis was exempt from human-subjects research regulations under 45 CFR 46.104. For all 140 wound specimens (93 internal, 47 external) data used in this study, paired swabs were collected from the same site during the same clinical encounter: one specimen was submitted for routine aerobic culture with AST and the other to the BioExcel Diagnostics laboratory for PCR testing. Thus, timing, collection site, and handling were equivalent across both index and reference tests, and turnaround differences reflected only laboratory workflow. As in our prior wound infection study, both PCR and culture were requested simultaneously by the treating provider as part of routine diagnostic practice.

### PCR Assays and Feature Derivation

To ensure standardized input for downstream logistic regression modeling, we utilized real-time PCR data previously generated in our comparative evaluation of PCR and culture for wound infections (9). In that study, bead-based mechanical lysis was used to disrupt both Gram-positive and Gram-negative organisms, followed by extraction with the MagMAX™ Microbiome Ultra Nucleic Acid Isolation Kit (Thermo Fisher Scientific). Amplification was performed on the SmartChip Real-Time PCR System (Takara Bio) using TaqMan probe–based assays (Thermo Fisher Scientific) targeting a broad panel of clinically relevant pathogens and antibiotic resistance genes. Each run incorporated internal controls for extraction efficiency, amplification performance, and contamination, along with positive and negative assay controls.

From this dataset, real-time PCR assays provided cycle-threshold (Ct) values for *S. aureus*, S*. lugdunensis*, coagulase-negative *staphylococci* (CoNS), S*. saprophyticus*, and *mecA*. Targets were considered positive if Ct ≤ 34. Raw Ct values for the *mecA* assay, *S. aureus*, and coagulase-negative *staphylococci* (CoNS) were recorded.

For each specimen, we calculated the gap between *mecA* and each organism as *dCt_S.aureus_ = (Ct_mecA_ - Ct_S.aureus_)* and *dCt_ConS_ = (Ct_mecA_ - Ct_Cons_).* When direction was not needed, the absolute output values for these equations (𝑎𝑑𝐶𝑡_𝑆.𝑎𝑢𝑟𝑒𝑢𝑠_ 𝑜𝑟 𝑎𝑑𝐶𝑡_𝐶𝑜𝑁𝑆_) were used. To quantify the closeness of *mecA* to each organism’s Ct value, we applied a logistic transform to the calculated absolute gaps using the equations below:

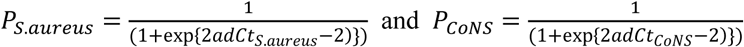

These P values were only defined when the corresponding Ct pair existed; otherwise, they were set to 0. ΔCt features were omitted if a component’s Ct was unavailable; sentinel Ct values (e.g., 40) were not employed for non-detects.

### Machine-Learning Model and Calibration

The primary classifier was logistic regression with class weighting (class_weight="balanced"). For linear models, we applied median imputation, followed by standardization and classification using scikit-learn pipelines. Median imputation was performed only on the training data. The pipeline included median imputation for missing values, feature standardization, and Platt probability calibration via “CalibratedClassifierCV (method = "sigmoid", cv=3)” inside each training split; the calibrated model then generated predictions for the held-out fold. Unless otherwise stated, decisions used a fixed threshold of 0.50 on calibrated probabilities. Five-fold stratified cross-validation generated out-of-fold (OOF) probability predictions (10–12). Model training and cross-validated evaluation were conducted in the 36 culture-confirmed *S. aureus*-positive cases (reference cohort). After cross-validation (CV), we fit the final calibrated model on all 36 specimens and generated predictions for the full 93-specimen dataset and eventually the external validation cohort. A biologic plausibility gate was applied post-prediction (never during CV or training). If PCR was negative for *S. aureus*, the MRSA probability was set to 0 and the qualitative output reported “No MSSA/MRSA.” When PCR was positive for *S. aureus*, we applied the 0.50 threshold to classify MRSA vs MSSA. External-cohort MRSA/MSSA attribution followed the same rule (classify only when PCR was positive for *S. aureus*).

Comparator models, linear SVM, random forest, and HistGradientBoosting, were trained on the same feature set with similar imputation methods; linear SVM used scaling and class weights, random forest used class_weight = "balanced_subsample," and HistGradientBoosting used only imputation. Class imbalance (25 MSSA versus 11 MRSA in the reference cohort) was addressed through stratified data splits and class weighting.

### Performance Metrics

Primary metrics that were evaluated are sensitivity, specificity, accuracy, ROC AUC, and Brier score, computed from OOF predictions. Calibration was assessed with reliability curves and expected calibration error (ECE). Uncertainty for performance estimates was summarized with non-parametric bootstrap resampling; additional robustness checks included per-fold summaries, 100×5 repeated cross-validation, and delete-1 jackknife influence (see Results). The Python code employed for the analysis is provided in the supplementary materials.

### Index Test and Blinding

All data outputs presented in this study were generated using a Python analysis pipeline (mrsa_audit_pipeline.py) without any manual adjustment, subjective interpretation, or retrospective reclassification of calls. For the internal cohort, although this was a retrospective dataset, performance metrics were calculated from out-of-fold predictions in cross-validation; thus, each case’s prediction was generated by a model that had no access to its own reference standard. For the external cohort, the trained model produced predictions without access to culture or AST results, which were only used for performance comparison.

## Results

### Dataset Composition

This study analyzed 93 wound and soft tissue infection specimens that were previously included in our comparative diagnostic evaluation of real-time PCR and culture for podiatric wound infections (9). In that study, PCR demonstrated 97% sensitivity and 95% specificity for detecting *S. aureus*, underscoring the robustness of the PCR workflow for detecting *S. aureus* in podiatric wound specimens (9). Within the present dataset (Table 1 and Supplementary Table S1), 38 of 93 specimens (40.9%) tested PCR-positive for *S. aureus*. Of these cases, 22 yielded coagulase-negative *Staphylococci* (CoNS) with overlapping Ct ranges, highlighting the challenge of accurately attributing *mecA* to *S. aureus* versus CoNS in polymicrobial wound specimens. Thirty-six specimens were culture-positive for *S. aureus*, comprising 25 methicillin-susceptible *S. aureus* (MSSA) and 11 MRSA. These 36 culture-confirmed cases were used as the reference cohort for model training and evaluation. The remaining 57 culture-negative specimens served as the test group for the biologic gating strategy, ensuring that the model was assessed in both *S. aureus*–positive and –negative contexts. This observed overlap between *S. aureus* and CoNS detections provided the rationale for deriving ΔCt-based features and developing a logistic regression model to assign *mecA* to its most likely s*taphylococcal* source.

**Table 1.**
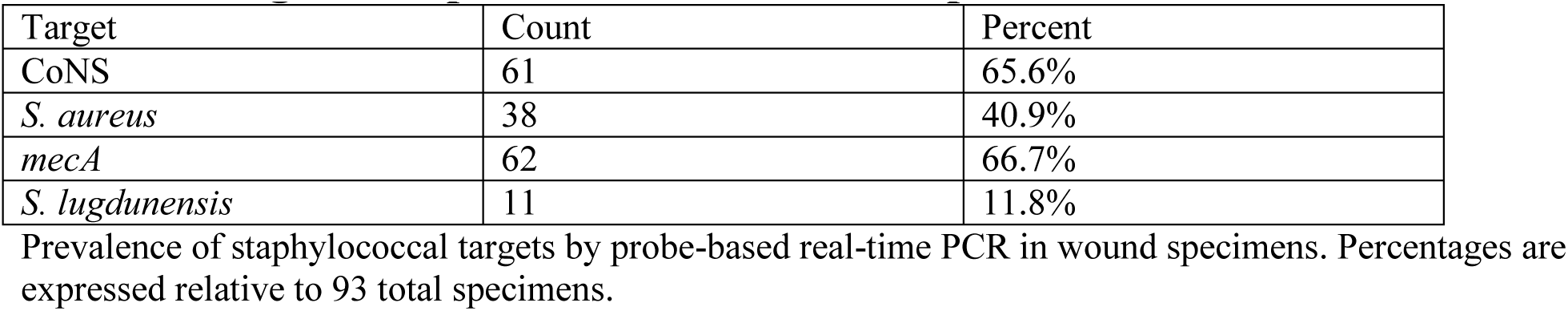
Target-level prevalence in 93 wound specimens.

### Comparator Model Performance

To benchmark performance across different machine learning approaches, we evaluated four classifiers using the same ΔCt-derived features and five-fold cross-validation: calibrated logistic regression, linear support vector machine (SVM), random forest, and histGradientBoosting (Table S2). Logistic regression (Figure 1A and 1B) achieved the best overall balance of discrimination and calibration, with an accuracy of 91.7%, an AUC of 0.931, and the lowest Brier score (0.093). The linear SVM performed similarly in terms of discrimination (AUC 0.927, accuracy 91.7%) but showed poorer probability calibration (Supplementary Figure S2A and S2B). Random forest performance was lower, with an AUC of 0.865 and an accuracy of 88.9% (Supplementary Figure S3A and S3B). HistGradientBoosting failed to generalize, with an AUC of 0.464 and accuracy of 69.4%, reflecting overfitting in this modest dataset (Supplementary Figure S4A and S4B). Among the four models, logistic regression provided the most balanced combination of discrimination and calibration, supporting its selection as the primary analytic framework for attributing *mecA* in polymicrobial wound specimens.

**Figure 1.**
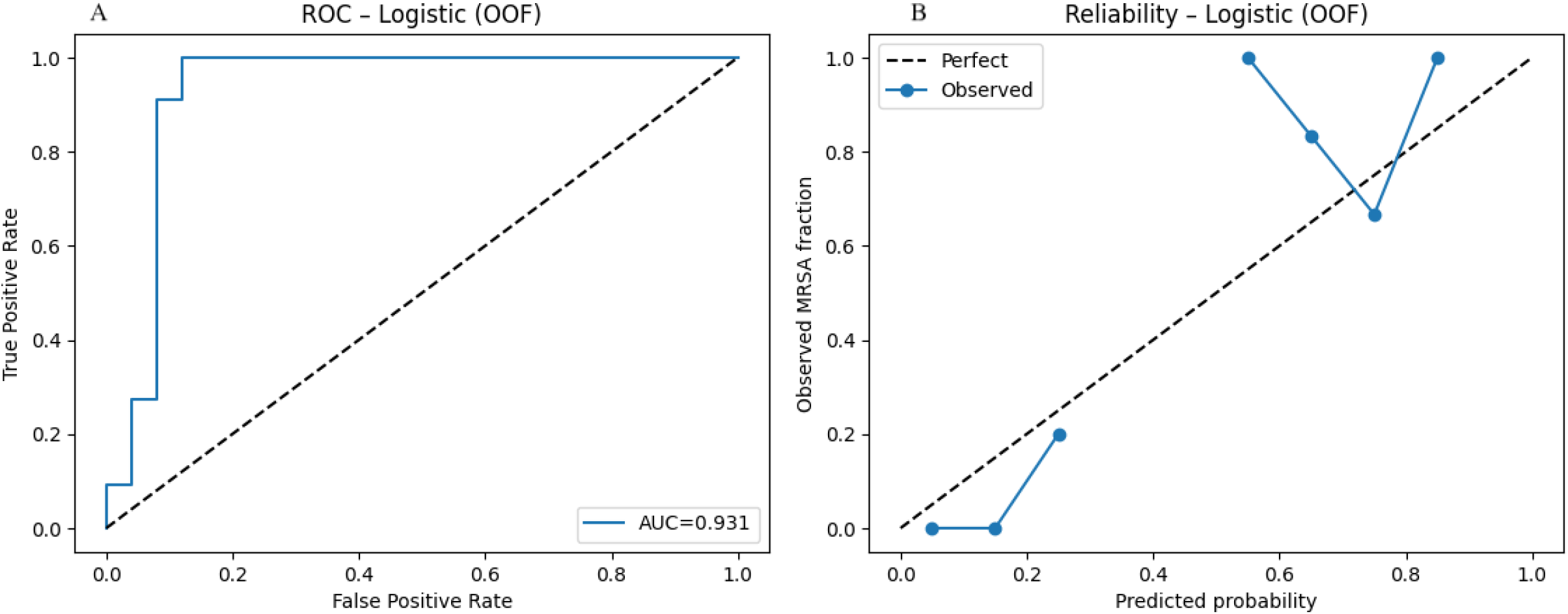
Logistic regression model’s discrimination and calibration. Receiver operating characteristic (ROC, panel A) and calibration (panel B) curves for logistic regression in the internal cohort (n = 93; 36 culture-confirmed S. aureus, including 11 MRSA). The model achieved an AUC of 0.931, accuracy of 91.7%, and a Brier score of 0.093, the lowest among all classifiers tested. ROC analysis confirmed strong discrimination, while calibration analysis showed probabilities were reasonably aligned with observed MRSA fractions, though slight underestimation occurred at intermediate probability levels.

### Logistic Regression Performance

In the 36 culture-confirmed S. aureus cases (Table 2), the logistic regression model achieved 91.7% overall accuracy (33/36). Sensitivity for MRSA was 90.9% (10/11), and specificity for MSSA was 92.0% (23/25). The cross-validated ROC AUC was 0.931, and the Brier score was 0.093 (Table 3). The confusion matrix contained 23 true negatives, 10 true positives, 2 false positives, and 1 false negative.

**Table 2.** Specimen provenance and PCR characteristics of 36 culture-positive Staphylococcus aureus cases.

| Case | Culture_SA | PCR_SA | Ct_SA | PCR_mecA | Ct_mecA | Ct_CoNS | dCt_SA | dCt_CoNS | AST_SA | fold | oof_prob | oof_pred | result |
| --- | --- | --- | --- | --- | --- | --- | --- | --- | --- | --- | --- | --- | --- |
| 3 | Positive | Positive | 19.15 | Negative | NaN | 33.29 | NaN | NaN | MSSA | 2 | 0.170304 | MSSA | TN |
| 6 | Positive | Negative | NaN | NaN | NaN | NaN | NaN | NaN | MRSA | 4 | 0.273535 | MSSA | FN |
| 10 | Positive | Positive | 28.90 | Negative | NaN | 31.93 | NaN | NaN | MSSA | 4 | 0.180230 | MSSA | TN |
| 12 | Positive | Positive | 19.18 | Positive | 20.24 | 22.37 | 1.06 | -2.13 | MSSA | 1 | 0.694728 | MRSA | FP |
| 15 | Positive | Positive | 16.17 | Positive | 32.50 | 21.60 | 16.33 | 10.90 | MSSA | 5 | 0.110147 | MSSA | TN |
| 16 | Positive | Positive | 19.17 | Positive | 25.05 | 27.60 | 5.88 | -2.55 | MSSA | 3 | 0.206905 | MSSA | TN |
| 17 | Positive | Positive | 17.14 | Negative | NaN | NaN | NaN | NaN | MSSA | 2 | 0.151834 | MSSA | TN |
| 19 | Positive | Positive | 15.34 | Positive | 14.57 | 33.96 | -0.77 | -19.39 | MRSA | 2 | 0.691481 | MRSA | TP |
| 22 | Positive | Positive | 17.14 | Positive | 28.18 | 28.43 | 11.04 | -0.25 | MSSA | 3 | 0.077272 | MSSA | TN |
| 25 | Positive | Positive | 24.35 | Positive | 25.03 | 28.47 | 0.68 | -3.44 | MRSA | 5 | 0.576307 | MRSA | TP |
| 27 | Positive | Positive | 18.80 | Positive | 18.02 | 24.00 | -0.78 | -5.98 | MRSA | 2 | 0.659835 | MRSA | TP |
| 28 | Positive | Positive | 18.70 | Positive | 33.77 | 27.50 | 15.07 | 6.27 | MSSA | 1 | 0.064823 | MSSA | TN |
| 36 | Positive | Positive | 19.02 | Positive | 31.16 | 30.80 | 12.14 | 0.36 | MSSA | 5 | 0.064932 | MSSA | TN |
| 39 | Positive | Positive | 17.02 | Negative | NaN | 28.00 | NaN | NaN | MSSA | 2 | 0.155345 | MSSA | TN |
| 41 | Positive | Positive | 20.02 | Positive | 30.51 | 33.94 | 10.49 | -3.43 | MSSA | 1 | 0.118248 | MSSA | TN |
| 45 | Positive | Positive | 19.02 | Positive | 33.96 | NaN | 14.94 | NaN | MSSA | 3 | 0.095549 | MSSA | TN |
| 48 | Positive | Positive | 20.50 | Positive | 19.75 | 33.00 | -0.75 | -13.25 | MRSA | 1 | 0.808624 | MRSA | TP |
| 55 | Positive | Positive | 22.94 | Positive | 32.87 | NaN | 9.93 | NaN | MSSA | 4 | 0.126855 | MSSA | TN |
| 56 | Positive | Positive | 20.85 | Positive | 33.37 | NaN | 12.52 | NaN | MSSA | 1 | 0.092327 | MSSA | TN |
| 58 | Positive | Positive | 21.85 | Positive | 21.02 | 32.70 | -0.83 | -11.68 | MSSA | 1 | 0.786034 | MRSA | FP |
| 71 | Positive | Positive | 18.70 | Positive | 17.91 | NaN | -0.79 | NaN | MRSA | 3 | 0.644986 | MRSA | TP |
| 75 | Positive | Positive | 18.62 | Negative | NaN | NaN | NaN | NaN | MSSA | 5 | 0.239502 | MSSA | TN |
| 79 | Positive | Positive | 21.79 | Positive | 24.95 | 28.17 | 3.16 | -3.22 | MSSA | 4 | 0.285449 | MSSA | TN |
| 83 | Positive | Positive | 21.44 | Positive | 20.53 | NaN | -0.91 | NaN | MRSA | 1 | 0.727802 | MRSA | TP |
| 84 | Positive | Positive | 22.40 | Positive | 21.11 | 31.32 | -1.29 | -10.21 | MRSA | 1 | 0.753935 | MRSA | TP |
| 86 | Positive | Positive | 24.70 | Negative | NaN | NaN | NaN | NaN | MSSA | 5 | 0.205249 | MSSA | TN |
| 88 | Positive | Positive | 26.40 | Positive | 25.12 | NaN | -1.28 | NaN | MRSA | 4 | 0.542250 | MRSA | TP |
| 89 | Positive | Positive | 20.70 | Positive | 20.03 | NaN | -0.67 | NaN | MRSA | 3 | 0.634048 | MRSA | TP |
| 92 | Positive | Positive | 27.00 | Positive | 33.49 | NaN | 6.49 | NaN | MSSA | 4 | 0.167130 | MSSA | TN |
| 93 | Positive | Positive | 30.10 | Negative | NaN | NaN | NaN | NaN | MSSA | 5 | 0.179034 | MSSA | TN |
| 95 | Positive | Positive | 22.15 | Positive | 33.58 | 34.14 | 11.43 | -0.56 | MSSA | 2 | 0.066046 | MSSA | TN |
| 98 | Positive | Positive | 26.45 | Positive | 30.90 | 31.83 | 4.45 | -0.93 | MSSA | 3 | 0.121615 | MSSA | TN |
| 100 | Positive | Positive | 31.30 | Negative | NaN | NaN | NaN | NaN | MSSA | 3 | 0.189011 | MSSA | TN |
| 103 | Positive | Positive | 16.90 | Positive | 15.89 | 20.42 | -1.01 | -4.53 | MRSA | 5 | 0.654238 | MRSA | TP |
| 104 | Positive | Positive | 30.70 | Negative | NaN | NaN | NaN | NaN | MSSA | 4 | 0.181366 | MSSA | TN |
| 107 | Positive | Positive | 16.12 | Positive | 25.37 | 25.89 | 9.25 | -0.52 | MSSA | 2 | 0.100909 | MSSA | TN |

Abbreviations: Ct\_SA – cycle threshold for *S. aureus*; Ct\_mecA – cycle threshold for *mecA*; Ct\_CoNS – cycle threshold for coagulase-negative *staphylococci*; dCt\_SA – Ct\_mecA minus Ct\_SA; dCt\_CoNS – Ct\_mecA minus Ct\_CoNS; AST\_SA – phenotypic susceptibility (MSSA vs MRSA); oof\_prob – out-of-fold predicted probability; oof\_pred – out-of-fold predicted label; result – classification outcome (TP, TN, FP, FN); MSSA – methicillin-sensitive *S. aureus*; MRSA – methicillin-resistant *S. aureus*; CoNS – coagulase-negative *staphylococci*; SA – *S. aureus*.

**Table 3.** Logistic regression performance (36 culture-positive cases, OOF CV)

| Metric | Estimate | 95% CI | Numerator/Denominator |
| --- | --- | --- | --- |
| Accuracy | 0.917 | 0.78–0.97 | 33/36 |
| Sensitivity | 0.909 | 0.62–0.98 | 10/11 |
| Specificity | 0.920 | 0.75–0.98 | 23/25 |
| Brier score | 0.093 | — | — |

Out-of-fold (OOF) cross-validation performance of the ΔCt-informed logistic regression in culture- positive *S. aureus* cases.

We also compared the model with a prevalence-only null model and calculated the Brier skill score (BSS), also known as the index of prediction accuracy (IPA). The model’s Brier score (0.093) was substantially lower than the null score (0.212, MRSA prevalence = 30.6%), corresponding to a BSS/IPA of 0.56, or a 56% improvement over baseline. Together with the AUC of 0.931, these results indicate strong discrimination and reliable probability estimates. Decision curve analysis further showed that the model provided greater net benefit than default strategies of treating all or treating none. At the prespecified 50% cutoff for out-of-fold probabilities, the model achieved a net benefit of 0.222, while the treat-all approach (i.e., with Anti-MRSA targeted antibiotics) produced a negative benefit (–0.389). These findings demonstrate that the model not only distinguishes MRSA from MSSA with high accuracy but also offers clear clinical value by reducing unnecessary MRSA therapy. Figure 2 shows the decision curve analysis across clinically relevant thresholds

**Figure 2.**
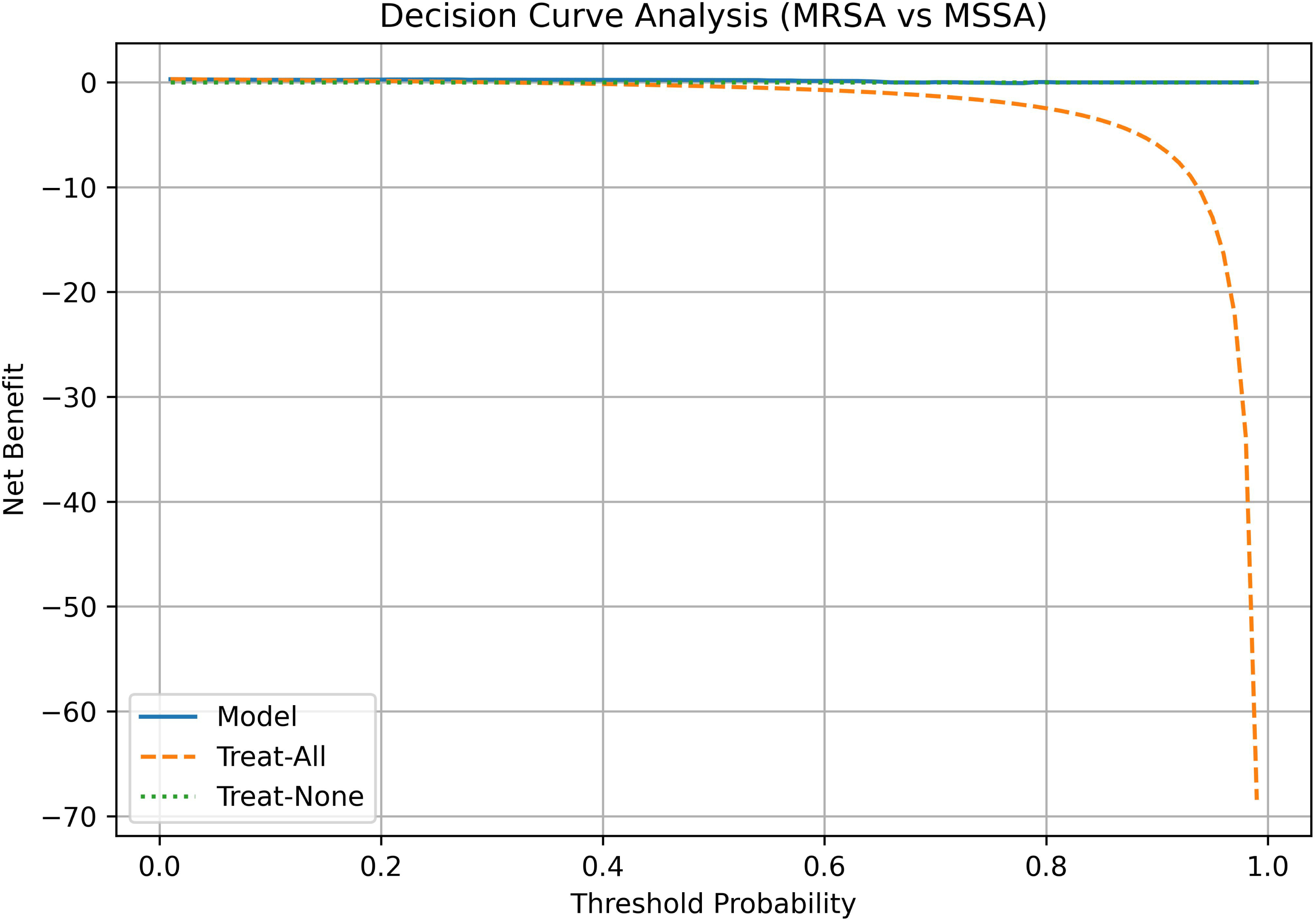
Decision curve analysis of the logistic regression model for MRSA versus MSSA. The logistic regression model showed consistently higher net benefit than default strategies of treating all or none across clinically relevant thresholds. At a 50% probability threshold, the model maintained a positive net benefit (0.222), whereas treat-all yielded a negative net benefit (–0.389).

### Model Calibration, Robustness, and Stability

Beyond point estimates of sensitivity and specificity, it is essential to evaluate whether a diagnostic model produces probabilities that are reliable, stable across resampling schemes, and robust to individual case influence. Therefore, we examined calibration, repeated cross-validation, fold-level performance, bootstrap resampling, and jackknife influence (Figures 3–7).

**Figure 3.**
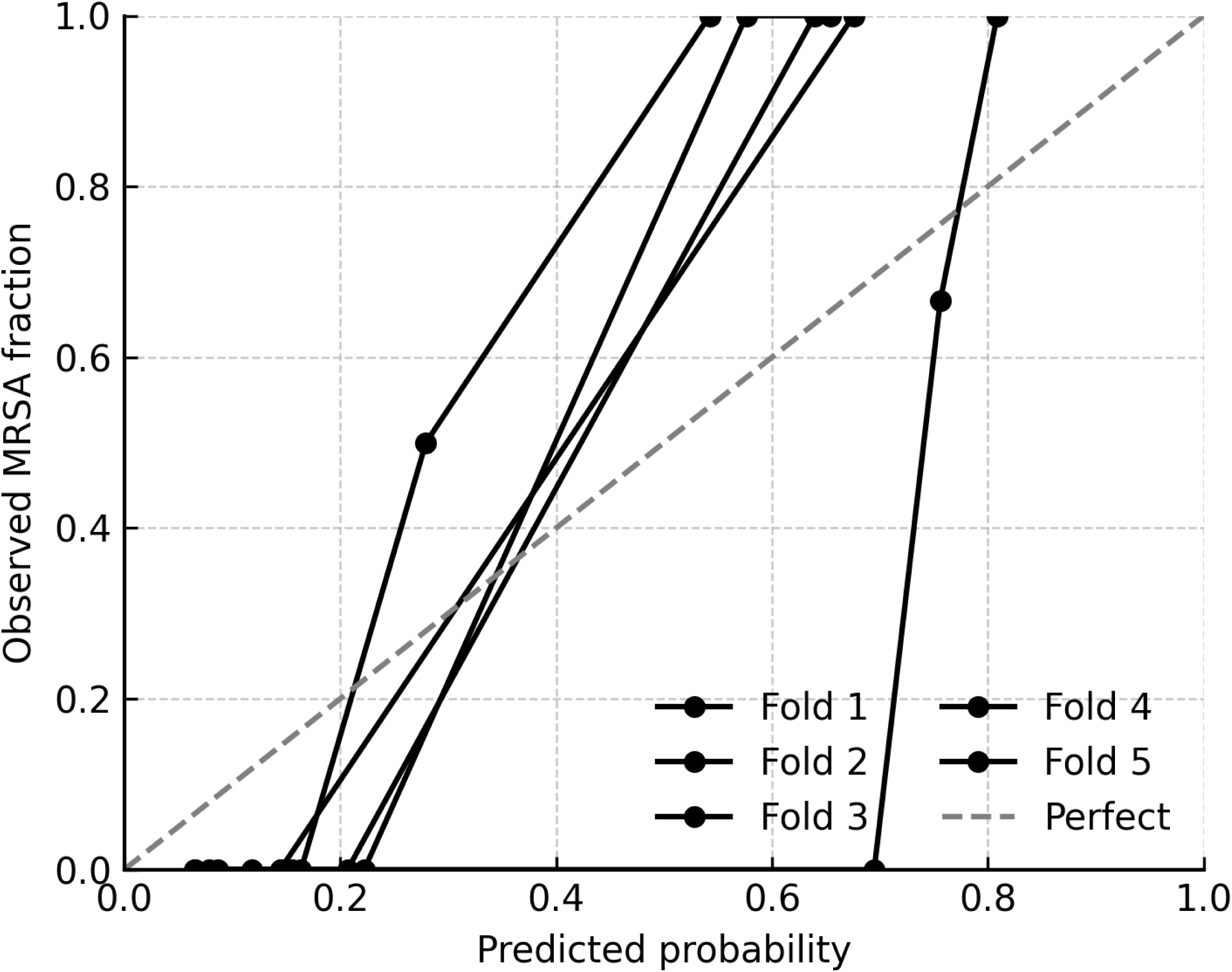
Calibration across five cross-validation folds. Calibration (reliability) curves are shown for each of the five cross-validation folds in the internal cohort. Curves deviated from the ideal 1:1 diagonal, particularly at mid-range predicted probabilities where the model underestimated observed MRSA prevalence. Expected calibration errors ranged from 0.18 to 0.22 (mean ≈ 0.20), consistent with the modest number of MRSA cases per fold. Despite moderate miscalibration, overall discrimination remained strong.

**Figure 4.**
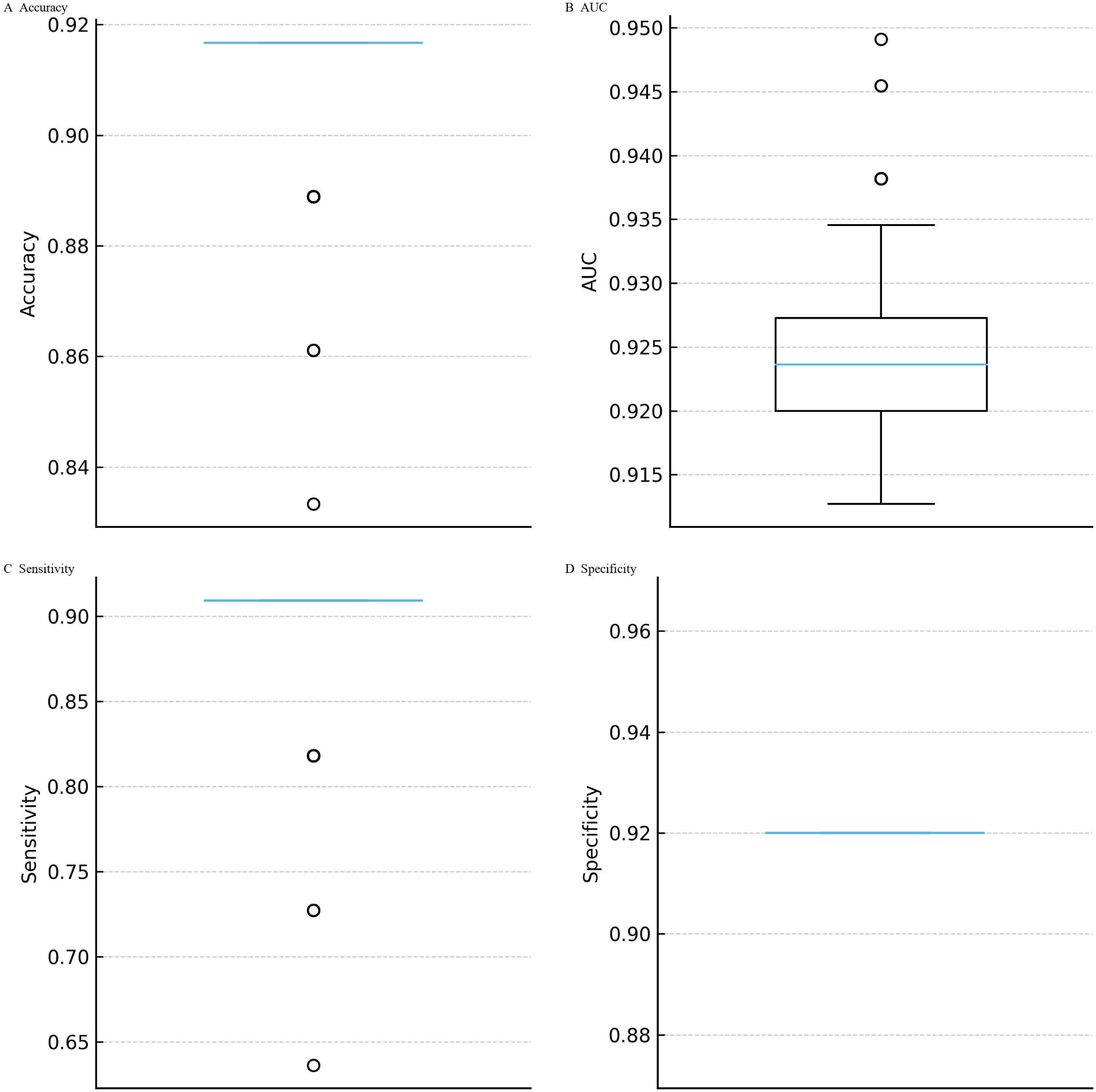
Stability of logistic regression performance across repeated cross-validation. Distributions of accuracy, AUC, sensitivity, and specificity are shown across 100 iterations of stratified 5-fold cross-validation. The mean accuracy was 0.911 (SD = 0.015; range 0.833– 0.917), mean AUC 0.926 (SD = 0.007; range 0.913–0.949), sensitivity 0.890 (SD = 0.049; range 0.636–0.909), and specificity consistently 0.920 (SD ≈ 0). Narrow interquartile ranges confirmed reproducibility of accuracy, AUC, and sensitivity. Variability was driven almost entirely by sensitivity, reflecting the limited number of MRSA cases per fold.

**Figure 5.**
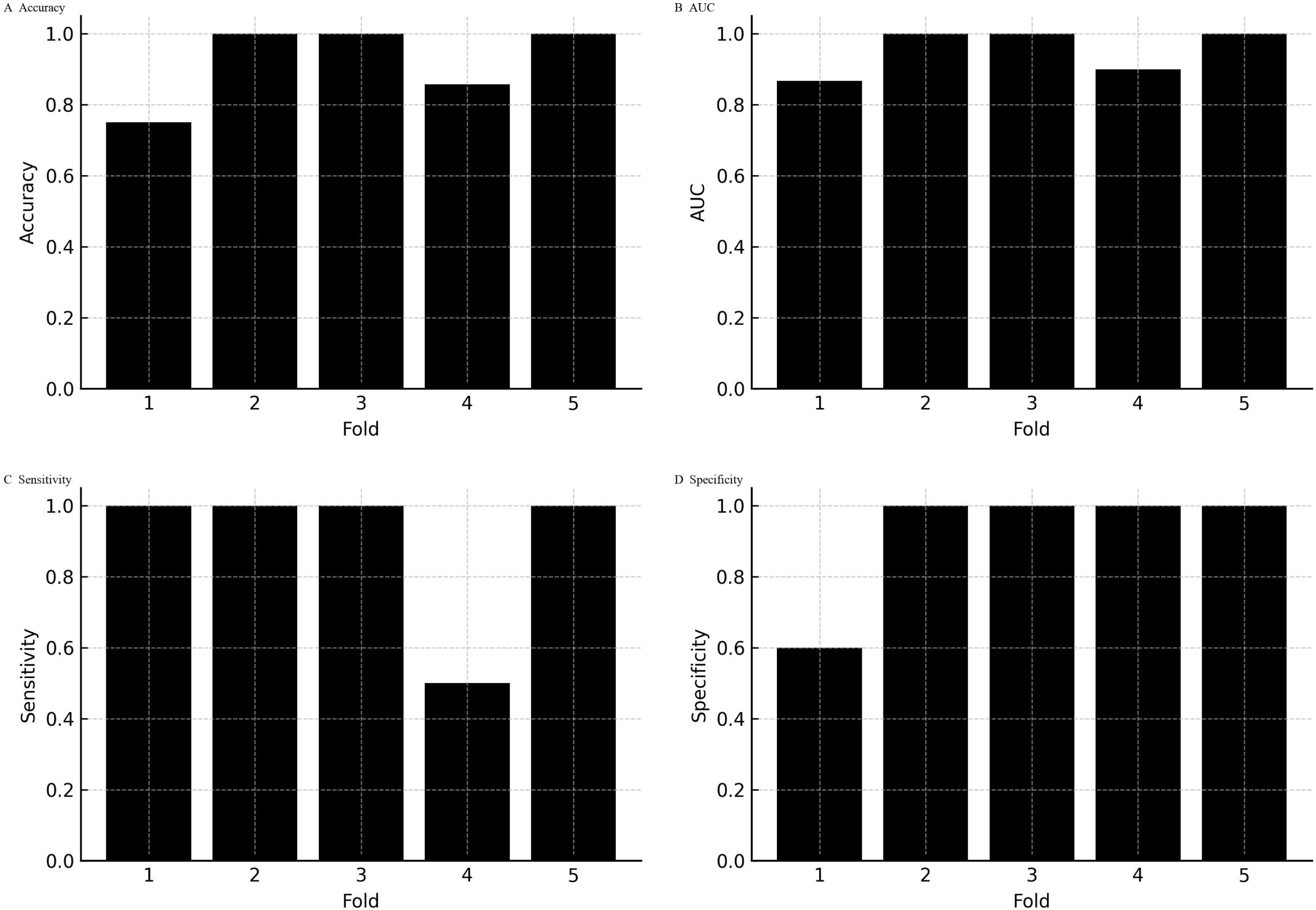
Per-fold performance within an exemplar split. Performance metrics are displayed for each of the five folds in a representative cross-validation split. Specificity ranged from 0.60 to 1.00 (mean 0.92; per-fold SD = 0.179), highlighting within-split variability despite consistently high average discrimination.

**Figure 6.**
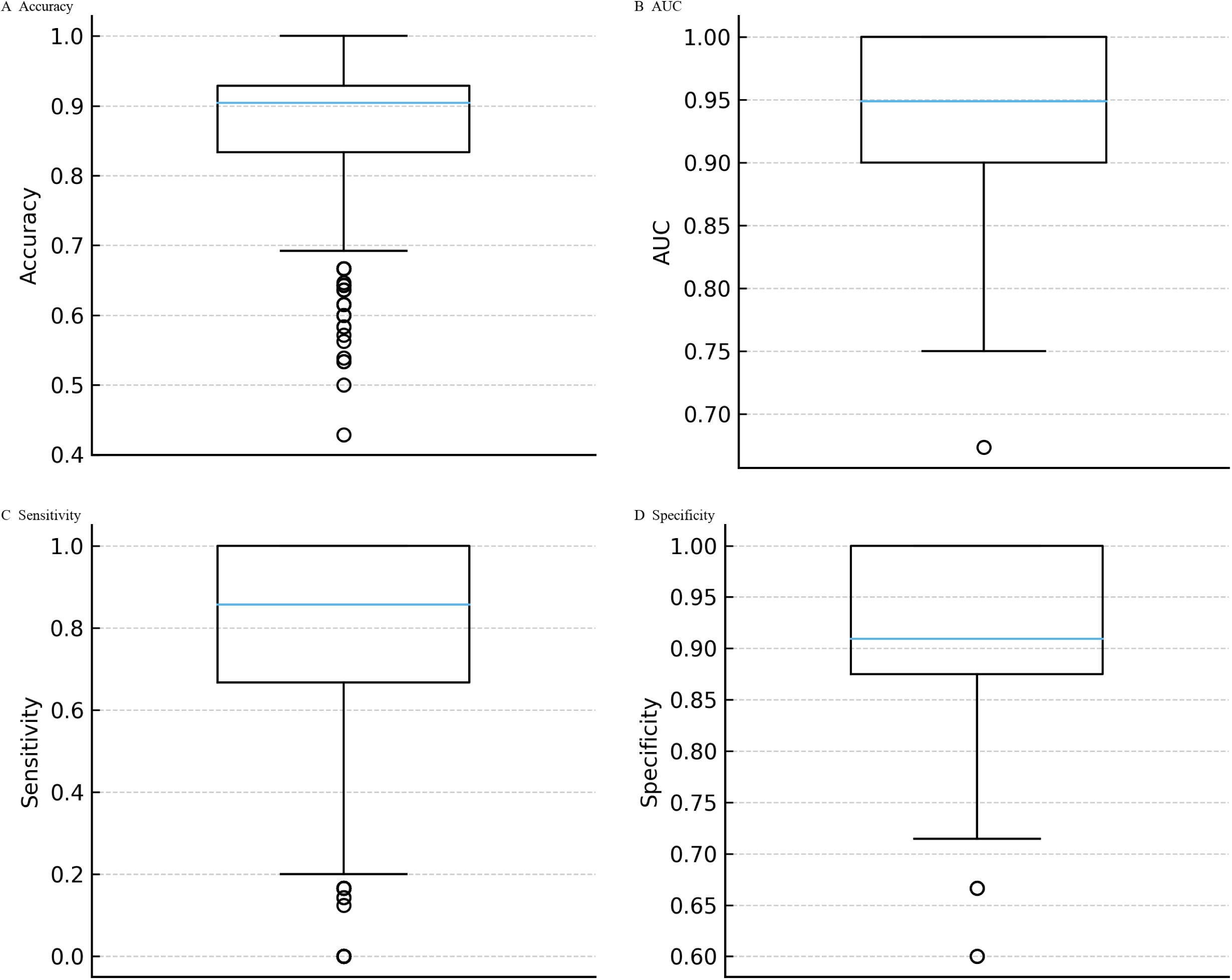
Bootstrap distributions of model performance. Nonparametric bootstrap resampling of the 36 culture-confirmed S. aureus cases (25 MSSA, 11 MRSA) was used to quantify uncertainty. Median estimates were 0.878 for accuracy (95% CI 0.64–1.00), 0.945 for AUC (0.83–1.00), 0.804 for sensitivity (0.17–1.00), and 0.921 for specificity (0.75–1.00). These distributions confirm that observed performance metrics are robust to resampling and not artefacts of a particular partitioning scheme.

**Figure 7.**
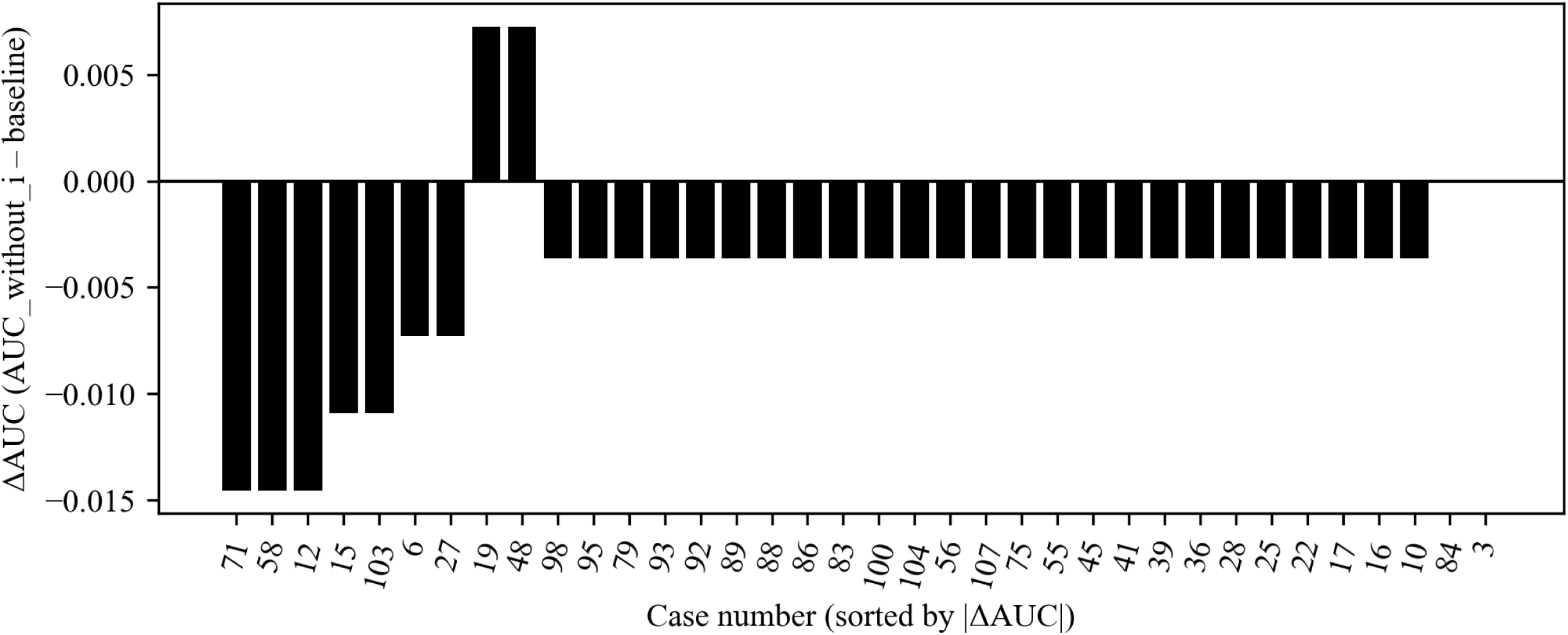
Jackknife validation of AUC estimates. Leave-one-out (jackknife) resampling of the 36 culture-confirmed S. aureus cases assessed the influence of individual specimens on AUC. Omitting any single case altered the AUC by no more than 0.015. Only five specimens produced absolute ΔAUC values > 0.01, each corresponding to borderline ΔCt values near the classification threshold. The remaining 31 specimens had negligible impact (|ΔAUC| ≤ 0.007), confirming the robustness of the logistic regression model to individual case influence.

Across the five cross-validation folds, the calibration curves deviated from the ideal 1:1 line, particularly at mid-range predicted probabilities where the model tended to underestimate the observed MRSA fraction. Expected calibration errors ranged from 0.18 to 0.22 (mean ≈ 0.20), reflecting moderate miscalibration consistent with the limited number of MRSA cases per fold (Figure 3). Despite this, classification metrics remained strong. Repeated 100×5-fold cross-validation demonstrated that the logistic regression model’s performance was stable across resampling runs. The mean accuracy was 0.911 (SD 0.015), with values ranging from 0.833 to 0.917. The mean AUC was 0.926 (SD 0.007), spanning 0.913 to 0.949. Sensitivity averaged 0.890 (SD 0.049), varying between 0.636 and 0.909, while specificity was consistently 0.920 across all folds (SD ≈0). Interquartile ranges were narrow for accuracy, AUC, and sensitivity, underscoring reproducibility of the model’s discrimination. Variability was almost entirely driven by sensitivity, reflecting the limited number of MRSA cases within each fold. (Figure 4). At the per-fold level within an exemplar split, specificity ranged 0.60–1.00 (mean 0.92; per-fold SD = 0.179), illustrating within-split variability (Figure 5). To summarize uncertainty, while avoiding aggregation ambiguity, nonparametric bootstrap over the 36 culture-confirmed cases gave specificity 0.921 (95% CI 0.75–1.00), sensitivity 0.804 (0.17–1.00), accuracy 0.878 (0.64–1.00), and AUC 0.945 (0.83–1.00) (Figure 6). These distributions confirm that the observed performance metrics are not artefacts of a single partitioning scheme. Finally, jackknife analysis showed that omitting any single case changed the AUC by no more than 0.015. Only five of the 36 specimens produced absolute ΔAUC values exceeding 0.01; these were cases with probabilities near the decision threshold, highlighting how borderline ΔCt values can slightly influence performance. The other 31 observations had negligible impact (|ΔAUC| ≤ 0.007), underscoring the robustness of the model to individual cases (Figure 7). Collectively, these analyses show that the logistic regression model generalizes well across resampling schemes and is robust to individual cases. However, its probability estimates exhibit moderate miscalibration and may require recalibration on larger cohorts.

### Application to the Full Cohort

The calibrated model with *S. aureus* gating was applied across all 93 wound specimens (Supplementary Table S3). For the 57 specimens that were culture-negative for *S. aureus*, the gating rule assigned an MRSA probability of zero and reported these as “No MSSA/MRSA.” As expected, none of these culture-negative wounds were misclassified as MRSA or MSSA, demonstrating that the gating strategy eliminated spurious resistant calls in the context of *mecA* detections arising from coagulase-negative staphylococci.

Among the 36 specimens that were culture-positive for S. aureus, the model produced 23 MSSA calls, 12 MRSA calls, and one “No MSSA/MRSA” assignment. This distribution aligns with the validated performance metrics observed in the AST-defined reference cohort (sensitivity 90.9% and specificity 92.0%), while extending the same biologic plausibility rules to the full dataset. Overall, model outputs across all 93 specimens consisted of 55 “No MSSA/MRSA,” 26 MSSA, and 12 MRSA classifications.

### External Validation

In the independent external cohort (n = 47), culture identified 13 specimens positive for *S. aureus*. The PCR panel detected *S. aureus* in 12 of these, yielding a sensitivity of 92.3% (12/13) compared with culture. Among 34 culture-negative specimens, PCR results were negative in 33 and positive in one, giving a specificity of 97.1% (33/34). This corresponded to one false-negative and one false-positive result relative to culture (Table 4). For methicillin resistance attribution, the biologic gating rule was applied so that only *S. aureus* PCR–positive specimens were classified as MRSA or MSSA. Within this subset (n = 12 evaluable specimens), the calibrated logistic regression model identified all MRSA cases correctly, corresponding to a sensitivity of 100% for MRSA detection. Among MSSA cases, one was misclassified as MRSA, giving a specificity of 87.5%. The overall accuracy for MRSA/MSSA classification in *S. aureus*–positive specimens was 91.7%, with positive and negative predictive values of 80% and 100%, respectively (Supplementary Table S4).

**Table 4.** Performance of PCR panel for *S. aureus* detection and MRSA attribution in internal and external cohorts.

| Task | Cohort | n evaluable | Sensitivity | Specificity | Accuracy | PPV | NPV |
| --- | --- | --- | --- | --- | --- | --- | --- |
| *S. aureus* detection vs culture | Internal (n=93) | 93 | 97.2% | 95.0% | 96.0% | 94.6% | 97.4% |
|  | External (n=47) | 47 | 92.3% | 97.1% | 95.7% | 92.3% | 97.1% |
| MRSA attribution (SA+ only) | Internal (n=36) | 36 | 90.9% | 92.0% | 91.7% | 83.3% | 95.8% |
|  | External (n=12) | 12 | 100% | 87.5% | 91.7% | 80.0% | 100% |

Notes: *S. aureus* (SA) detection was evaluated against culture as the reference. MRSA attribution was evaluated only among *S. aureus* PCR-positive specimens, after applying the biologic gating rule. PPV and NPV are influenced by cohort prevalence.

## Discussion

Polymicrobial wound specimens frequently contain *S. aureus* alongside coagulase-negative staphylococci (CoNS), many of which carry the *mecA* gene. This overlap complicates interpretation, because *mecA* co-detection with *S. aureus* by PCR does not necessarily imply methicillin-resistant *S. aureus* (MRSA). Misattribution of *mecA* to *S. aureus* may result in unnecessary use of glycopeptides or oxazolidinones rather than the recommended β-lactams for MSSA, exposing patients to greater toxicity, higher cost, and more complicated treatment courses. (1, 2).

To address this challenge, we incorporated ΔCt features, defined as the Ct difference between *mecA* and staphylococcal targets such as *S. aureus* or CoNS, into a calibrated, class-weighted logistic regression model, combined with a biologic gate that suppressed MRSA/MSSA classification when *S. aureus* PCR was negative. In the 36 culture-confirmed S. aureus cases, our logistic regression model achieved 91.7% accuracy, with sensitivity for MRSA of 90.9% and specificity for MSSA of 92.0%. The model showed strong discrimination (AUC = 0.931) and a low Brier score (0.093), indicating overall good calibration and discrimination. However, because the Brier score is prevalence-dependent and does not directly capture clinical value (13), we also compared the model with a prevalence-only null model. This yielded a Brier skill score of 0.56, reflecting a 56% improvement in probabilistic accuracy over baseline. To further evaluate clinical usefulness, we applied decision curve analysis. At the prespecified 50% probability cutoff, the model achieved a net benefit of 0.222, while the treat-all approach, equivalent to giving MRSA-directed or considered antibiotics to every *S. aureus* case, resulted in a negative net benefit (–0.389). These findings show that the model not only distinguishes MRSA from MSSA accurately but also guides more rational antibiotic choices by reducing unnecessary MRSA therapy.

Our approach builds on prior ΔCt-based strategies used in commercial assays (6). For example, the MRSA/SA ELITe MGB kit interprets ΔCt <2 cycles between *mecA* and an *S. aureus* target as MRSA, with 100% concordance in a study of 82 nasal isolates, whereas the Xpert MRSA/SA assay showed lower concordance (76.8%) (1, 2, 6). A key limitation of these approaches is that they do not explicitly account for coagulase-negative staphylococci (CoNS), which frequently harbor *mecA* and are common in wound specimens. This omission increases the risk of misclassifying *mecA*-positive CoNS as MRSA. Our model achieved comparable accuracy, 91.7% in the external cohort and 95.7% in the internal cohorts, while providing probabilistic outputs and explicit biologic gating. Unlike fixed cutoffs, logistic regression incorporates multiple features (Ct, ΔCt, detection flags) and generates calibrated probabilities that can be adjusted to clinical settings.

Selective broth enrichment combined with *nuc*–*mecA* PCR has also been proposed, achieving 93.5% sensitivity and 88.6% specificity in 1,250 samples (7). Our sensitivity (90.9%) and specificity (92.0%) fall within this range, but our method operates directly on wound specimens that are often polymicrobial without enrichment. Third-generation droplet digital PCR (ddPCR) assays have demonstrated excellent performance (>96% sensitivity, >91% specificity) in nasal swabs (14), but require specialized instrumentation and still risk CoNS-driven misclassification. Our framework achieved comparable accuracy in wound specimens using probe-based real-time PCR assays, while incorporating the linkage probability to CoNS. Similarly, rapid multiplex PCR (15) and MALDI-TOF combined with machine learning (16) show the breadth of approaches for MRSA detection, yet differ in instrumentation requirements and scope. Our ΔCt- informed logistic regression complements these efforts by focusing on direct clinical wound specimens and explicitly linking *mecA* attribution to biologic plausibility. Our work extends this concept by embedding ΔCt features within a machine-learning framework, calibrating the resulting probabilities, and applying biologic gating to eliminate implausible MRSA calls.

In our wound cohort, 25 of 36 culture-positive S. aureus cases were methicillin-sensitive (MSSA; 69%), underscoring the importance of distinguishing between *S. aureus* and CoNS as carriers of mecA. A similar distribution was observed in the external validation set, where 8 of 12 *S. aureus*–positive cases were MSSA (67%) and 4 were MRSA (33%). By combining ΔCt quantification with biologic gating, our framework directly addresses this limitation, reduces overtreatment with MRSA-directed agents, and supports appropriate β-lactam use for MSSA. The use of calibrated probabilities further provides transparency and auditability, features that may enhance clinician confidence in molecular resistance reporting. While stand-alone MRSA PCR kits exist and are valuable for screening, they are not designed for polymicrobial wound specimen testing. Our framework is intended to enhance the panel-based TaqMan assays already offered as LDTs in CAP/CLIA labs, improving mecA attribution within multiplex workflows without requiring additional testing.

As a secondary finding, PCR demonstrated high accuracy for *S. aureus* detection in the external validation cohort (n = 47), with 92.3% sensitivity, 97.1% specificity, and 95.7% overall accuracy, closely matching the performance observed in our prior evaluation of the internal cohort.

### Limitations and Future Directions

The main limitation of this study was the modest sample size, particularly the small number of MRSA cases, which contributed to variability in sensitivity estimates and limited generalizability. Culture served as the reference standard but may misclassify borderline or mixed infections. Although the underlying classification performance was strong, as shown by AUC values, Brier score, and positive net benefit, the reliability curves show moderate miscalibration. Larger external cohorts are needed to confirm performance, refine calibration, and quantify inter-laboratory variability.

## Conclusions

Our ΔCt-informed, *S. aureus*–gated logistic regression provided accurate and interpretable *mecA* attribution in wound specimens, with balanced sensitivity and specificity and fewer implausible MRSA calls. By embedding biologic plausibility and PCR kinetics, the model reduced overtreatment with MRSA-directed antibiotics. Importantly, decision curve analysis demonstrated that this approach yielded a positive net benefit compared with treating all *S. aureus* cases as MRSA, indicating that the model enhances both clinical decision-making and statistical accuracy. PCR also maintained high accuracy for S. aureus detection in an external validation set, underscoring the robustness of the molecular workflow. Although our findings are encouraging, the number of MRSA cases was modest, and larger, more diverse cohorts will be needed to validate calibration, refine cutoff rules, and confirm clinical utility. If validated, this framework could be integrated into standard probe-based wound PCR panel workflows to improve antibiotic stewardship and decrease the use of unnecessary anti-MRSA agents in complex infections.

## Data and Code Availability

All analysis artifacts, including the case-level provenance tables with out-of-fold probabilities and fold assignments, the 93-case calibrated model outputs with *S. aureus* PCR gating applied and the reproducible Python script used for analysis (mrsa_audit_pipeline.py), are available from the authors upon reasonable request in accordance with institutional data-sharing policies.

## Ethics Statement

This study analyzed de-identified data originally generated for routine diagnostic purposes. No new specimens were collected, and the secondary analysis meets criteria for exemption under 45 CFR 46.104.

## Author Contributions

M.D. conceived the study. M.D. oversaw study design, data analysis, and manuscript preparation. H.N., S. D., contributed to data collection, analysis, interpretation, and critical manuscript review. H.L.M. and K.H.L. provided clinical oversight and critical manuscript review.

## Declaration of Generative AI and AI-Assisted Technologies in the Writing Process

During preparation of this manuscript, the authors used ChatGPT (OpenAI) only for language polishing and formatting (e.g., grammar, phrasing, and consistency of tense). No protected health information or proprietary data were entered into the tool. All scientific content (study design, data analysis, interpretation, and conclusions) was written and verified by the authors. The authors reviewed and edited all AI-assisted text and accept full responsibility for the manuscript’s content.

